# Embryonic loss of human females with partial trisomy 19 identifies region critical for the single active X

**DOI:** 10.1101/119495

**Authors:** Barbara R Migeon, Michael A Beer, Hans T Bjornsson

**Affiliations:** McKusick Nathans Institute of Genetic Medicine, Baltimore MD; USA; Department of Pediatrics, Johns Hopkins University, School of Medicine, Baltimore MD; USA; Department of Biomedical Engineering, Johns Hopkins University School of Medicine, Baltimore MD; USA

## Abstract

To compensate for the sex difference in the number of X chromosomes, human females, like human males have only one active X. The other X chromosomes in cells of both sexes are silenced *in utero* by *XIST*, the *Inactive X Specific Transcript gene*, that is present on all X chromosomes. To investigate the means by which the human active X is protected from silencing by *XIST*, we updated the search for a key dosage sensitive *XIST* repressor using new cytogenetic data with more precise resolution. Here, based on a previously unknown sex bias in copy number variations, we identify a unique region in our genome, and propose candidate genes that lie within, as they could inactivate *XIST*. Unlike males, the females who duplicate this region of chromosome 19 (partial 19 trisomy) do not survive embryogenesis; this preimplantation loss of females may be one reason that more human males are born than females.

## Introduction

The concept of a single active X was introduced by Mary Lyon in her 1962 paper [1], wherein she extended her hypothesis from mice to other mammals, especially humans. She pointed out that a single X is sufficient for survival (i.e., Turner syndrome) and that no matter the number of X chromosomes in both sexes, only one was active (i.e., human sex chromosome aneuploidies, 47,XXY, 49,XXXXY)). The developmental pathway leading to X dosage compensation is not limited to human females, nor is it inhibited by a Y-chromosome [2, 3]. In fact, no compelling evidence precludes the likelihood that it maintains the activity of the single X chromosome in normal males as well as females.

Initially, the term *single active X* was used as a synonym for X inactivation [4, 5]. However, it might be more appropriate to refer to Lyon’s hypothesis as the single active X, rather than the X inactivation hypothesis [6]. Although inactive X chromosomes are created in the process, they may not be the targets of the events that *initiate* dosage compensation of the human X [6].

The difference between the X *inactivation* and the *single active X* hypotheses is whether the underlying mechanisms count X chromosomes to determine how many should be inactive—that is, *choose the inactive X*, or alternatively, they *choose the single active X–* that is, the only fully functional X chromosome in each cell. Either mechanism is plausible, yet evidence based on studies of normal and aneuploid humans favors the single active X hypothesis, at least in our species:

*XIST* has all the hallmarks of a housekeeping gene: no TATA box, ubiquitous expression, and a 5’ CpG island that is methylated in inactive genes; furthermore, *XIST* is expressed from all X-chromosomes in the human zygote of either sex [7] albeit at low levels, until the time in embryogenesis when the locus on the future active X is turned off, and its CpG island becomes methylated in both males and females [8].

In addition, studies of 69, XXX and 69, XXY triploid cells provide compelling evidence that it is the *active* X that is chosen [9]. In contrast to 47,XXY and 47,XXX diploid cells that have a single active X, the majority of human triploid cells (87% of the 47 triploids studied) have two active X chromosomes [9-15]. This suggests that the extra set of autosomes in triploid cells allows the majority of these cells to maintain the activity of more than one X chromosome.

The simplest explanation for two active X chromosomes in triploid cells is that active X’s are chosen by repressing their *XIST* loci; the key repressor is encoded by an autosome, and the extra dose of this autosome and therefore of this key repressor leads to more than one active X [6, 9]. This *XIST repressor hypothesis* is depicted in Fig 1.

**Fig 1.**
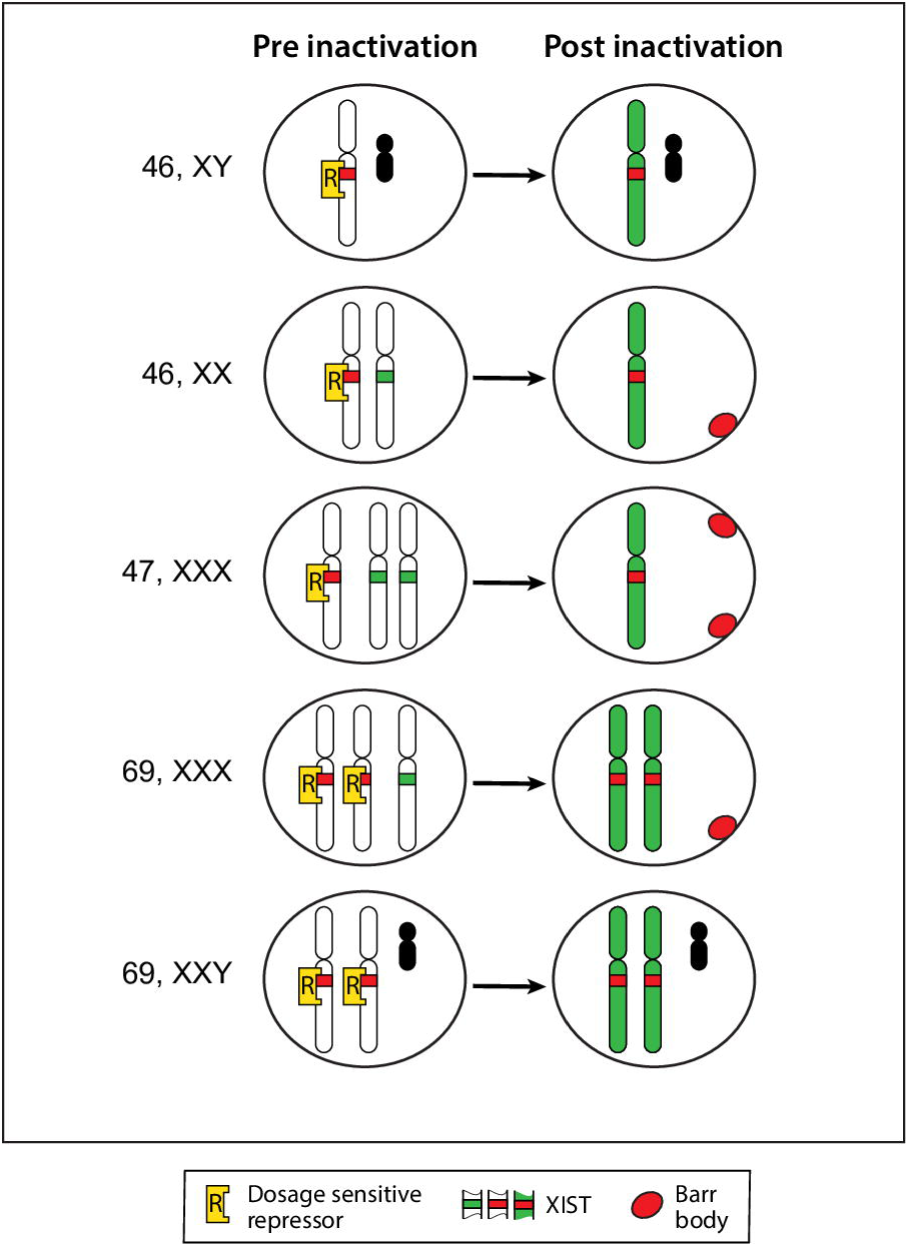
The XIST repressor model for the single active X. Our model depicts the putative dosage sensitive repressor(s) (yellow), which inactivate *XIST*, thereby protecting one X chromosome from inactivation in diploid 46,XX, 46,XY and 47,XXX cells, thus-directly choosing the active X (green). The non-blocked X chromosomes are inactivated by *XIST* transcription, becoming Barr bodies (red). In triploid cells (69,XXX, 69,XXY), more than one X is active because of the extra amount of the putative *XIST* repressor, contributed by the extra set of autosomes. The Y chromosome is depicted (black).

Clearly, the findings in triploids have implications for diploid cells; the most direct way to designate an active X in any cell would be to turn off its *XIST* locus. In male as well as female diploid cells – irrespective of the number of X’s in the cell – repression of *XIST* on one of them would insure an active X. All other X chromosomes would be silenced because their non-repressed *XIST* locus is subject to up-regulation. Therefore, by default, any chromosome with an active *XIST* gene will be silenced by the cascade of events induced by its transcripts [16].

The diploid: triploid difference in the number of active X’s points to a dosage sensitive autosomal gene, capable of turning off *XIST* on future active X chromosomes. In the case of triploidy, the triple dose of that repressor permits two X‘s to remain active in the majority of cells. If there were three copies of that repressor in *diploid* cells (i.e., in trisomies), we expect that both the X chromosomes in that cell might be active in many cells – presumably a very early lethal event, as two intact active X’s have never been observed, even among aborted fetuses.

The strategy for finding the chromosome that carried the putative *XIST* repressor was as follows: The X chromosome, itself, could be eliminated as the candidate chromosome because 47, triple X females have only a single active X (i.e. Fig 2b in Migeon et al. [17]). To identify the relevant autosome, Jacobs et al. [12] and Migeon et al. [9, 17] looked for two active X chromosomes in diploid cells with three copies of a single autosome. These studies of autosomal trisomies, surviving long enough to be recognizable conceptuses, eliminate 20 of the 22 autosomes as repressors of human *XIST*, based on the presence of a single *inactive* X in all the trisomic cells analyzed.

However, two full-trisomies, chromosomes 1 and 19, were never observed among miscarriages, presumably because triplication of these gene-dense chromosomes does not let the embryo survive long enough to be identified as a miscarriage. There-fore, for chromosomes 1 and 19 we could only study the partial trisomies that survive gestation. We expected such partially trisomic survivors would help us eliminate regions of the chromosome, associated with normal X inactivation, whereas the regions, never seen in live-born individuals, might encode the dosage sensitive *XIST* repressors. Migeon et al. [9] identified triplication of regions of chromosomes 1 and 19, associated with post implantation survival and normal patterns of X inactivation, and based on the relatively crude estimates of cytogenetic breakpoints available at the time, excluded them as candidate regions for the *XIST* repressor. But, as females with inherited triplications of 1p31, 1p21.3–q25.3, or 19p13.2–q13.33 were never observed, we suggested that these regions might contain key dose-sensitive gene(s) that induce *XIST* repression.

Since these exclusion studies based on cytogenetic analysis were reported in 2008 [9], extensive data have been collected on copy number variation in human patients; now that breakpoints of partial trisomies are definable at the level of base pairs, it seems appropriate to update the candidate regions on chromosomes 1 and 19. Revisiting this project has allowed us to find the autosome we were looking for, and to identify candidate genes. It has also revealed a previously unknown sex bias in copy number variants unique to one autosome. This sex bias implies a substantial pre-implantation loss of human females.

Our strategy for identifying suitable candidates for the key *XIST* repressor on chromosomes 1 and 19 was based on the following assumptions:

1. There were epigenetic factors that could repress *XIST in trans*. Of particular interest were the writers and erasers of epigenetic marks as lysine demethylases have been implicated in *Xist* activation in mice. In elegant experiments, Fukuda et al [18] showed that Kdm4b turns off the second active X in female mouse parthenotes by turning on their *Xist* locus. In addition, when the *Kdm2b* gene is deleted in female mice, investigators observed upregulation of *Xist*, which *decreased* the expression of X-linked genes, and dysregulated the expression of autosomal genes by affecting the protein complex that mediates *Xist* silencing [19]. If such lysine demethylases could activate *Xist* expression, then we speculated that other epigenetic marks could prevent it, perhaps by histone changes leading to DNA methylation.
2. Duplication of such a gene either by translocation or *in* situ amplification could turn off *XIST* on both X chromosomes in females leading to two active X chromosomes, a lethal event; however, males with their single X are not at risk of having more than a single active X chromosome. Therefore, candidate genes should show sex differences in the frequency of duplications: partial trisomies in males, but not in females.
3. Duplications that were *de novo* variants might have arisen after X inactivation was initiated, and therefore would not be expected to influence the event.
4. Deletions of candidate genes on one of a pair of autosomes would not be a problem in either sex, as the product of a single allele could be sufficient to turn off a single *XIST* locus in both sexes.

Therefore, based on these assumptions, we expected to see duplications only in males but deletions in both sexes.

## Results

### Chromosome 1 and 19 are rich in epigenetic players

Our search of the OMIM library and the UCSC genome browser revealed that many genes on our candidate chromosomes were known to have a role in transcriptional activation and repression. We learned that the three key proteins required for *Xist* silencing [20] were located on human chromosome 1: the lamin B receptor (LBR), SPEN (SHARP), and HNRNPU (SAFA) (Table 1). In addition, other relevant chromosome 1 genes include those encoding RBM15, HDAC1, SETDB1, the YY1 associated protein, YY1AP, the *XIST* activator KDM4A and potential *XIST* repressors KDM1A and KDM5B, which remove the transcriptional activator, H3K4. Similarly, chromosome 19 (Table 2) has the *Xist* activator KDM4B [18] and transcription activator (KMT2B), several potential *Xist* repressors, including the maintenance DNA methyltransferase, (DNMT1), the protein that tethers DNMT1 to chromatin and represses retrotransposons and imprinted genes (UHRF1) [21], the twin scaffold attachment factors SAFB & B2, the histone deacetylase, (SIRT6), a long non-coding RNA (TINCR) and clusters of zinc finger proteins, among others. We will show that human chromosome 1 plays a role in silencing X chromosomes, and human chromosome 19 seems to play a role in protecting the active X.

**Table 1.** **Genes on Chromosomes 1, showing total copy number variants, duplications and deletions, according to sex (M male, F female).** Genes shown on chromosome 1 are those known to have a role in silencing the inactive X as well as those that could play a role in silencing XIST. The 5’BP is based on GRCh38 when available or from the DECIPHER database (GRCh37). *Genes within previous candidate region [9].

| CHROM | GENE | 5'BP | Variants |  | Duplications |  | Deletions |  | Function |
| --- | --- | --- | --- | --- | --- | --- | --- | --- | --- |
|  |  |  | F | M | F | M | F | M |  |
| 1q42.12 | LBR | 225,401,501 | 14 | 4 | 6 | 3 | 1 | 1 | Lamin B Receptor |
| 1q44 | HNRNPU | 244,842,122 | 39 | 26 | 12 | 10 | 8 | 6 | Scaffold attachment factor A; *Xist* activator |
| 1p36.13 | SPEN | 16,174,359 | 3 | 5 | 0 | 1 | 2 | 1 | Transcriptional repressor |
| 1p35.2 | HDAC1 | 32,292,106 | 3 | 1 | 0 | 1 | 0 | 0 | Histone deacetylase |
| *1p13.3 | RBM15 | 110,338,425 | 6 | 7 | 1 | 0 | 5 | 2 | Member of Spen family. |
| *1q24.2 | METTL18 | 169,792,528 | 11 | 6 | 1 | 0 | 5 | 6 | Methylates non-histone proteins |
| 1p36.21 | PRDM2 | 13,698,874 | 6 | 10 | 0 | 1 | 6 | 5 | ZNF regulator of transcription |
| 1p36.12 | HP1BP3 | 20,742,676 | 4 | 4 | 0 | 2 | 0 | 1 | H3.2 (in all tissues) |
| *1q22 | YY1AP1 | 155,659,441 | 7 | 8 | 2 | 3 | 1 | 1 | YY1 associated protein |
| *1q25.3 | RNF2 | 185,014,551 | 6 | 4 | 1 | 0 | 3 | 3 | Represses imprinted genes |
| 1p36.31 | CHD5 | 6,101,786 | 24 | 17 | 0 | 1 | 12 | 11 | Swi/Snf p53 associated |
| *1q21.3 | SETDB1 | 150,515,756 | 3 | 4 | 0 | 2 | 0 | 0 | Methylates H3K9; |
| 1p34.2 | KDM4A | 44,115,796 | 3 | 1 | 0 | 0 | 1 | 1 | Possible Xist activator |
| 1p36.2 | KDM1A | 23,019,442 | 3 | 3 | 0 | 1 | 2 | 0 | Represses H3K4 |
| 1q32.1 | KDM5B | 202,727,402 | 5 | 6 | 2 | 4 | 1 | 1 | Represses H3K4 |
| *1p31.1 | ZNF265 | 71,063,290 | 9 | 6 | 1 | 2 | 5 | 2 | Ubiquitous zinc finger protein |
| *1p31.3 | MIER1 | 66,924,894 | 7 | 5 | 1 | 3 | 2 | 1 | Transcription repressor |
| *1p31.3 | TRIM33 | 114,392,776 | 7 | 8 | 1 | 2 | 4 | 3 | Represses transcription by binding to TSSs |
| *1q22 | ASHIL | 155,335,260 | 7 | 7 | 2 | 4 | 1 | 0 | Lysine methyltransferase 2H |
| *1q22 | OCT1 | 167,220,828 | 6 | 6 | 0 | 0 | 4 | 5 | Transcription factor in early zygote |

**Table 2.**
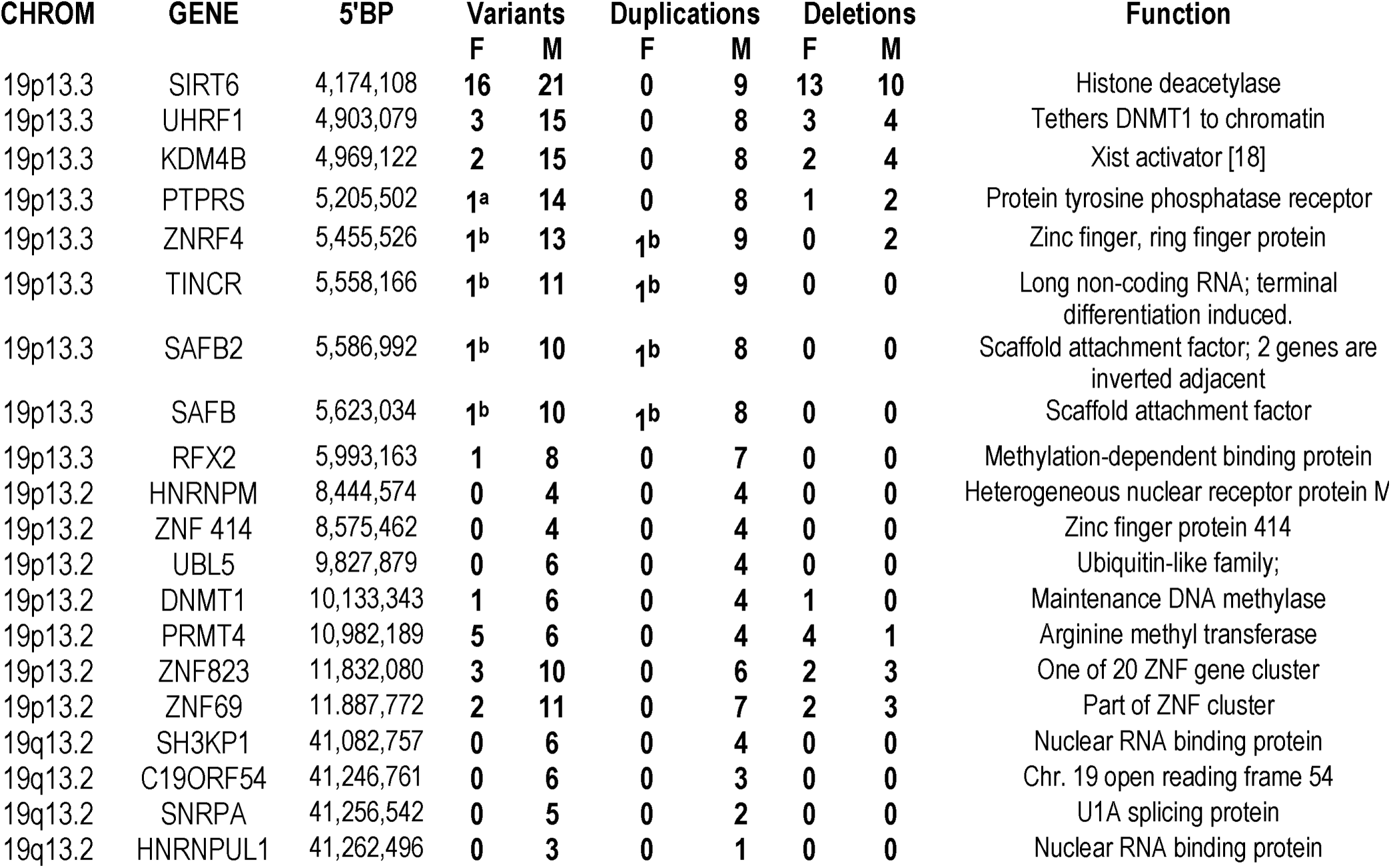
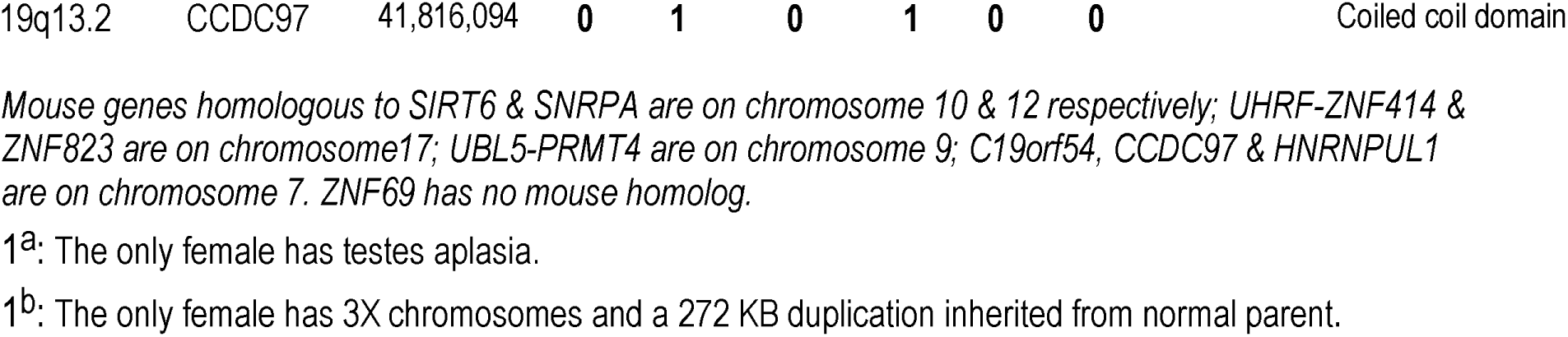
**Candidate genes on chromosome 19**, **showing total variants, duplications and deletions, according to sex (F female, M male).** The 5’BP is based on GRCh38 when available or from the DECIPHER database (GRCh37). These genes are all within the region of interest [9].

| CHROM | GENE | 5'BP | Variants |  | Duplications |  | Deletions |  | Function |
| --- | --- | --- | --- | --- | --- | --- | --- | --- | --- |
|  |  |  | F | M | F | M | F | M |  |
| 19p13.3 | SIRT6 | 4,174,108 | 16 | 21 | 0 | 9 | 13 | 10 | Histone deacetylase |
| 19p13.3 | UHRF1 | 4,903,079 | 3 | 15 | 0 | 8 | 3 | 4 | Tethers DNMT1 to chromatin |
| 19p13.3 | KDM4B | 4,969,122 | 2 | 15 | 0 | 8 | 2 | 4 | Xist activator [18] |
| 19p13.3 | PTPRS | 5,205,502 | 1<sup>a</sup> | 14 | 0 | 8 | 1 | 2 | Protein tyrosine phosphatase receptor |
| 19p13.3 | ZNRF4 | 5,455,526 | 1<sup>b</sup> | 13 | 1<sup>b</sup> | 9 | 0 | 2 | Zinc finger, ring finger protein |
| 19p13.3 | TINCR | 5,558,166 | 1<sup>b</sup> | 11 | 1<sup>b</sup> | 9 | 0 | 0 | Long non-coding RNA; terminal differentiation induced. |
| 19p13.3 | SAFB2 | 5,586,992 | 1<sup>b</sup> | 10 | 1<sup>b</sup> | 8 | 0 | 0 | Scaffold attachment factor; 2 genes are inverted adjacent |
| 19p13.3 | SAFB | 5,623,034 | 1<sup>b</sup> | 10 | 1<sup>b</sup> | 8 | 0 | 0 | Scaffold attachment factor |
| 19p13.3 | RFX2 | 5,993,163 | 1 | 8 | 0 | 7 | 0 | 0 | Methylation-dependent binding protein |
| 19p13.2 | HNRNPM | 8,444,574 | 0 | 4 | 0 | 4 | 0 | 0 | Heterogeneous nuclear receptor protein M |
| 19p13.2 | ZNF 414 | 8,575,462 | 0 | 4 | 0 | 4 | 0 | 0 | Zinc finger protein 414 |
| 19p13.2 | UBL5 | 9,827,879 | 0 | 6 | 0 | 4 | 0 | 0 | Ubiquitin-like family; |
| 19p13.2 | DNMT1 | 10,133,343 | 1 | 6 | 0 | 4 | 1 | 0 | Maintenance DNA methylase |
| 19p13.2 | PRMT4 | 10,982,189 | 5 | 6 | 0 | 4 | 4 | 1 | Arginine methyl transferase |
| 19p13.2 | ZNF823 | 11,832,080 | 3 | 10 | 0 | 6 | 2 | 3 | One of 20 ZNF gene cluster |
| 19p13.2 | ZNF69 | 11,887,772 | 2 | 11 | 0 | 7 | 2 | 3 | Part of ZNF cluster |
| 19q13.2 | SH3KP1 | 41,082,757 | 0 | 6 | 0 | 4 | 0 | 0 | Nuclear RNA binding protein |
| 19q13.2 | C19ORF54 | 41,246,761 | 0 | 6 | 0 | 3 | 0 | 0 | Chr. 19 open reading frame 54 |
| 19q13.2 | SNRPA | 41,256,542 | 0 | 5 | 0 | 2 | 0 | 0 | U1A splicing protein |
| 19q13.2 | HNRNPUL1 | 41,262,496 | 0 | 3 | 0 | 1 | 0 | 0 | Nuclear RNA binding protein |
| 19q13.2 | CCDC97 | 41,816,094 | 0 | 1 | 0 | 1 | 0 | 0 | Coiled coil domain |

*Mouse genes homologous to SIRT6 & SNRPA are on chromosome 10 & 12 respectively; UHRF-ZNF414 & ZNF823 are on chromosome 17; UBL5-PRMT4 are on chromosome 9; C19orf54, CCDC97 & HNRNPUL1 are on chromosome 7. ZNF69 has no mouse homolog.*
1<sup>a</sup>: The only female has testes aplasia.
1<sup>b</sup>: The only female has 3X chromosomes and a 272 KB duplication inherited from normal parent.

### Sex specific differences in variants for some epigenetic factors

Our search of the DECIPHER database that reports chromosomal gains (gene duplications and partial duplications) as well as losses (gene deletions and partial gene deletions) revealed the following:

Without exception, for all the epigenetic factors surveyed, we observed copy number variants in individuals who, although often clinically abnormal, survived gestation. However, in some cases, there were gains, but no losses, and in others, vice versa. As shown in Tables 1 and 2, many variants were equally distributed among males and females; however, some duplications showed marked sex specificity (Table 2).

None of the genes for epigenetic factors that we examined on chromosome 1 showed *extensive* sex specific differences in incidence of copy number variation (Table 1). However, four genes on chromosome 19 had the pattern we were looking for (Table 2): *KDM4B* (MIM 609765): two females vs. fifteen males had copy number variants. Of the fifteen male variants, eight were duplications and four were deletions; there were no female duplications and the two females had heterozygous deletions. Of the possible *XIST* repressors, there were three genes with similar patterns: 1) *DNMT1* (MIM 126375): a single female vs. six males had variants; four of the six males had duplications; the single female had a heterozygous deletion of the gene. 2). *UHRF1* (MIM 607990): of the three females vs. fifteen males with variants, no female vs. eight males had duplications; all three females had heterozygous deletions, and 3). *HNRNPM* (MIM 106994), the heterogeneous nuclear receptor protein: no variants in females, and four males with duplications.

These observations led us to systematically explore the sex bias along chromosome 1 and 19 by plotting a marker of this bias, the posterior gain rate of selected individual genes, in serial order from short arm telomere (*pter*) to the long arm telomere (*qter*) using gains as the copy number variant for the screen (See Methods and Figs 2A and 2B). Our screen revealed regions involving three cytogenetic bands on chromosome 19, in which the sex ratio of gains was skewed (19p13.3, 19p13.2 and 19q13.2) (Fig 2A); No-extensive sex bias was seen in candidate regions of chromosome 1 (Fig 2B).

**Fig 2.**
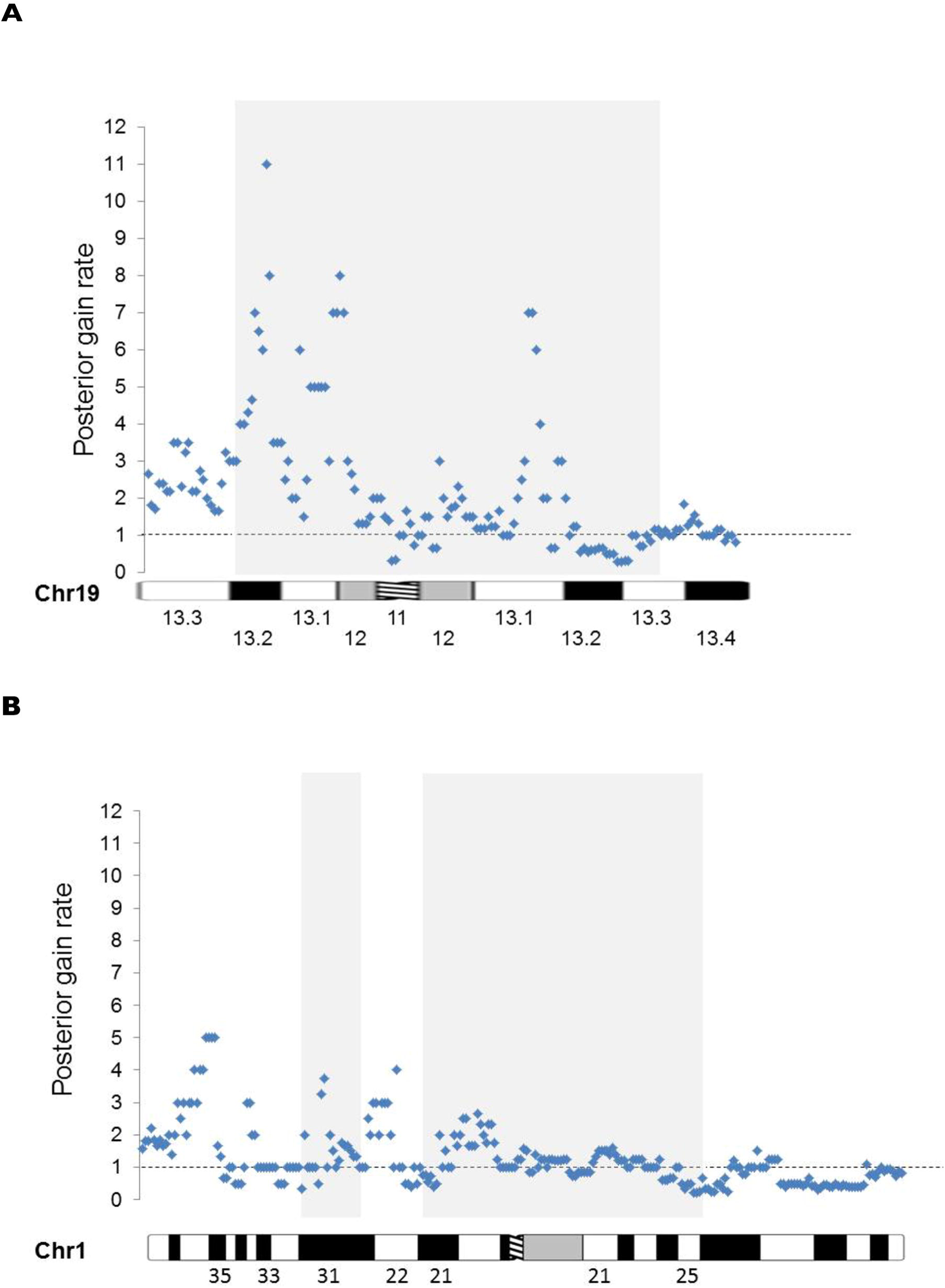
Systematic skewing of posterior gain rate (M:F sex ratio) on chromosome 19 (A) but not chromosome 1 (B). Because the lack of gains in either males or females leads to a zero value in either the nominator or denominator of a male/female gain ratio we calculated a posterior rate, which accurately reflects any observed deviation from the expected equal amount of male/female gains(=1). The grey box shows previous candidate regions, based on living females with partial trisomies. Note abnormal M:F posterior gain ratio in the candidate region of chromosome 19, but not on chromosome 1.

An expanded search at 500KB intervals throughout the 59 MB of chromosome 19 showed sex skewing not only in the numbers of duplications, but also in the total copy number variants, throughout an eight MB region of the short arm with tapering of skewing upstream and downstream of this region (Fig 3, Tables 2, 3, and S1 Table, showing details of the analysis, depicted in Fig 3). Within this ∼ eight MB region (from *PLIN4-ZNF709: 4*,*502*,*179 – 12*,*484*,*829bp* (*GRch38*), involving at least 237 genes and 1732 individuals with copy number variants) we observed a consistent loss of total variants in females, and absence of females with inherited duplications (See Figs 3A and 3B). Rarely, there was a female with a *de novo* duplication that because of its size, affected several genes in the region; such *de novo* variants may have arisen after X inactivation had occurred. We also observed a second notable region on the long arm of chromosome 19: Within 41 and 41.5 MB along the chromosome (with approximately 22 genes), there is a less striking loss of female variants, specifically duplications (See Figs 2B, 3A and 3B). Of note, both domains are within the candidate region reported by Migeon et al, 2008, [9] (Fig 2B, grey boxes).

**Fig 3.**
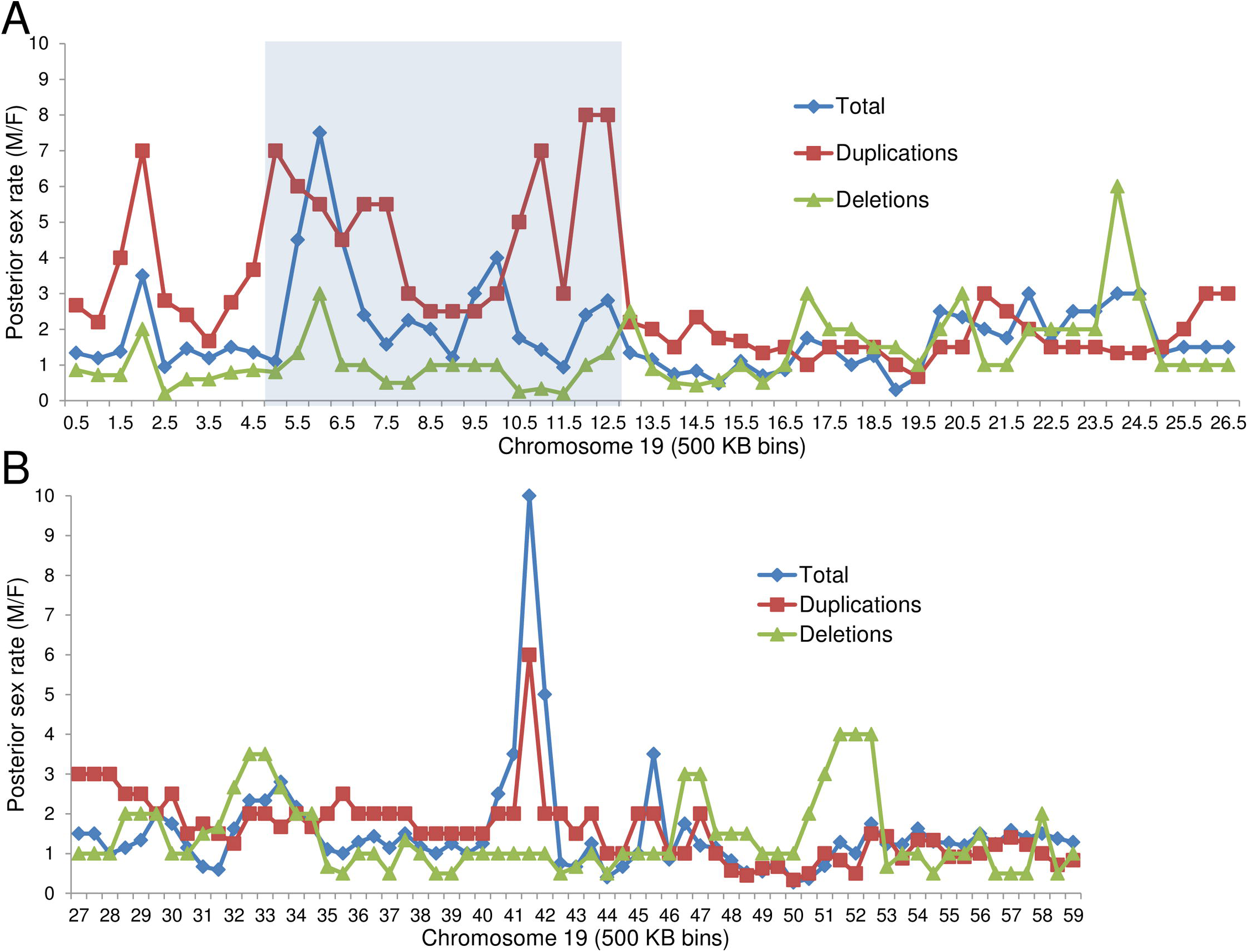
Posterior rate (M:F sex ratio) of duplications, deletions and total variants on chromosome 19, at 500 kb intervals throughout the p (A) and q (B) arms of the chromosome from pter to qter. Because the lack of gains in either males or females leads to a zero value in either the nominator or denominator of a male/female gain ratio we calculated a posterior rate, which accurately reflects any observed deviation from the expected equal amount of male/female gains(=1). Note the excess of males for both total variants and duplications in the domain 4.5 -12.5 MB on the short arm (A) as well as a smaller peak from 41.0-41.5 MB on the long arm (B) of chromosome 19.

To estimate the significance of the observed sex differences in duplications on chromosome 19, we used a permutation test.

We distributed the observed 852 total duplications on chromosome 19 randomly by sex and at random positions in 500KB bins of that chromosome. We then calculated the largest average M/F posterior sex ratio in a 8MB window observed along the chromosome. In the actual data, the highlighted 8MB region from bin 4.5MB to 12.5MB has an average M/F posterior sex ratio of 4.833. After 10^7^ random simulations, the highest observed average M/F posterior ratio for duplications was 3.52. Thus, the simulations directly show that the significance of a ratio of 4.833 is far less than p=10^-7^. We can estimate how many more simulations would be required to observe a ratio of 4.833 by plotting the distribution of the observed maximum average ratio in the simulations, as shown in S2 Fig. The tail of this distribution is approximately power law with exponent m^-9^, where m is the maximum average ratio, so we estimate that the simulations would have to be run 10^4^ or 10^5^ times longer to observe an average posterior sex ratio greater than the observed value of 4.833 on chromosome 19 once by chance, putting a conservative upper bound on p-value of 10^-11^.

### Extensive loss of females with partial trisomy is unique to chromosome 19

We looked for sex differences on all the other human autosomes using total gains (duplications and partial duplications) as the determinant for this screen (Figs A-T in S1 Fig.). Although the total gains in some genes originated more from males than females (i.e., Figs F and K in S1 Fig showing chromosomes 7 and 12), in others, more gains originated from females than males (See Figs N, O and Q in S1 Fig, showing chromosomes 15, 16 and 18). No sector showed the extensive domain of skewing, observed on the short arm of chromosome 19. The exceptional regions on chromosome 7 and 12 were assayed again, this time, with respect to duplications; we found that in every case they were limited to relatively few genes – unlike that seen for the chromosome 19 short arm. In addition, our previous studies showed that full trisomies 7 and 12 have a single active X [12]. We suggest that regions with skewed sex ratios on other chromosomes merit future study to determine the reason for the more localized sex differences.

### Zinc Finger Protein Clusters

On chromosome 19 there are several dense clusters of zinc finger proteins that differ with respect to sex ratio of variants and female loss (i.e., See Tables 2 and 3). These dense clusters of zinc finger proteins are known to be unique to chromosome 19 [22] and our survey of several other chromosomes attest to their uniqueness on chromosome 19. They are thought to be repressors of endogenous retroviruses (ERV) [22], which have been implicated in providing novel transcription factor binding sites, and generating novel functional lncRNAs [23]. One of the dense clusters is in the middle of 19p13.2, and this is the only cluster that shows the skewed sex ratio for total variants and duplications (Table 3).

**Table 3.**
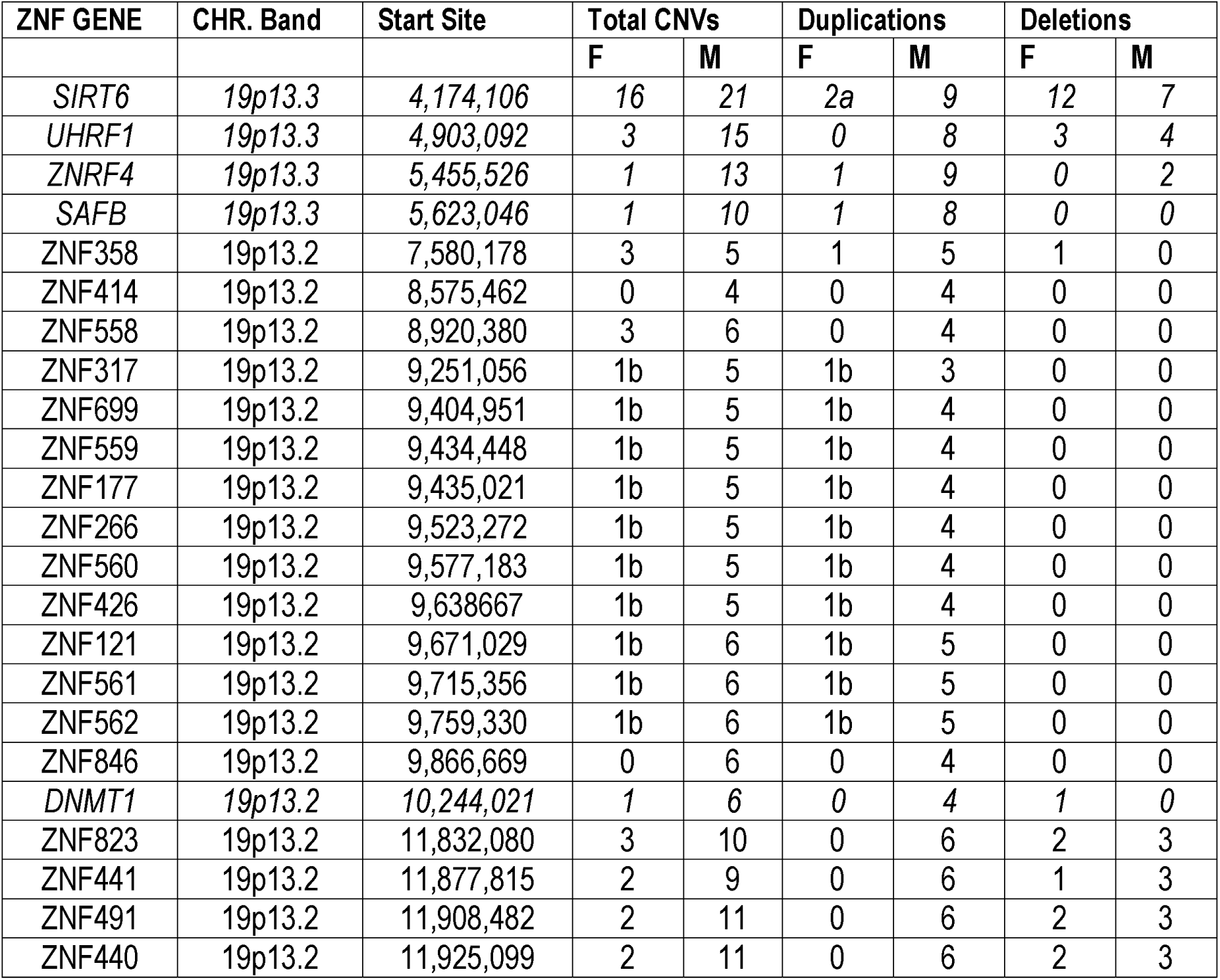
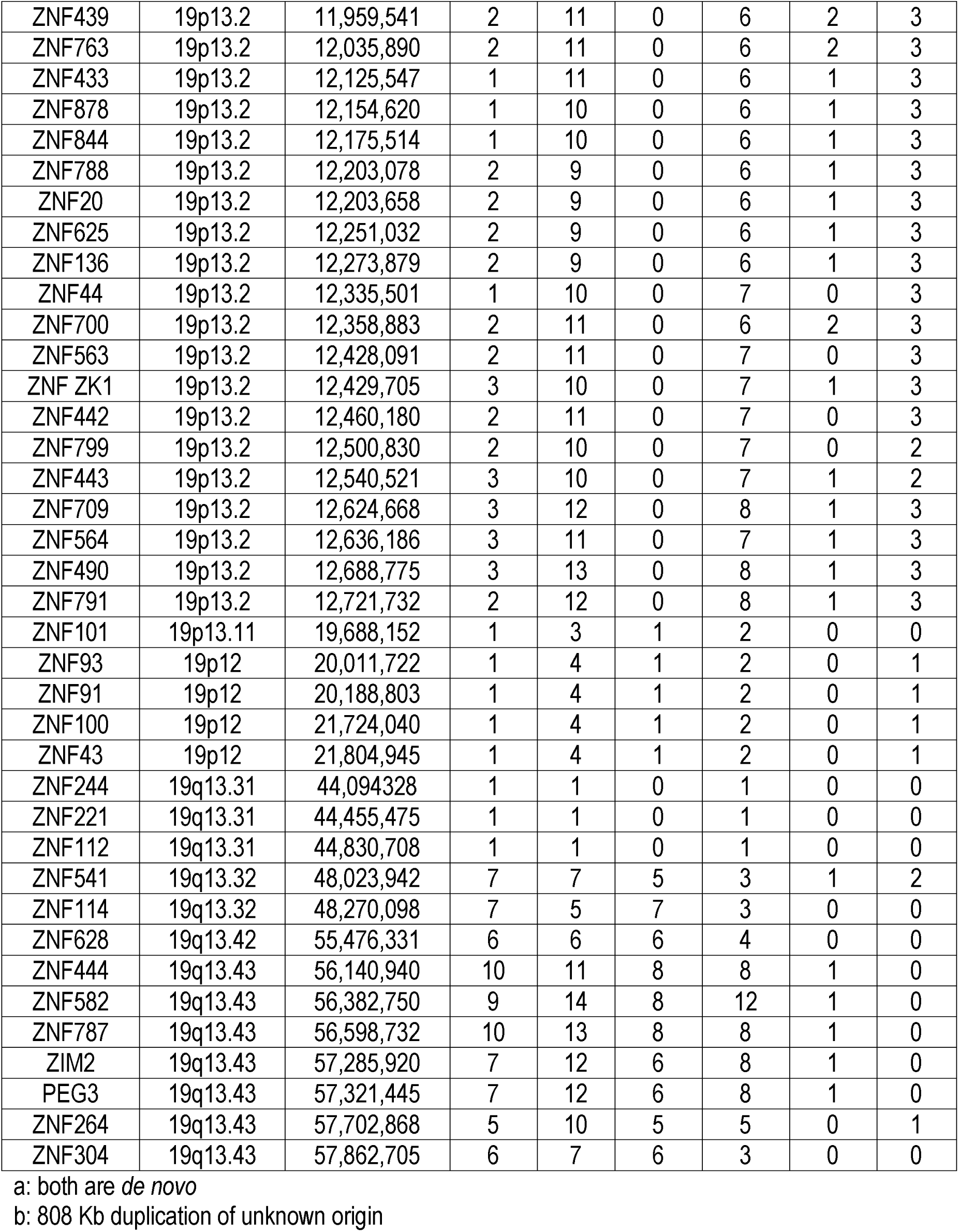
**Analysis of chromosome 19 clusters of zinc finger genes and their distribution**. Not all of the zinc finger genes on the long arm are shown, as their patterns are not remarkable. Also shown are the locations of some candidate genes (in italics) with respect to zinc finger clusters.

Another zinc finger protein REX1 (ZFP42, MIM 614572) is a known *Xist* repressor in the mouse [24]. Our observations indicate the need to further investigate the role of the chromosome 19p13.2 clusters of zinc finger proteins in maintaining the single active X.

### Gene content within exceptional regions of chromosome 19

Table 2 shows the candidate genes located within the 4.5–12.5 MB and 41.0–41.5 MB regions of interest. There are also unknown open reading frames, as well as many genes that are unlikely to be related to X inactivation, including those for the LDL receptor and TGFB1. Except for zinc finger genes and unknown open reading frames, we could not identify obvious candidates among the 22 genes in the 41-41.5 MB region of 19q13, but among them, we noted the splicing factor SNRPA with no female variants vs. five male variants (*XIST* & DNMT1 have spiced variants), and a conserved open reading frame locus (C19 orf54) with no female variants vs. six males variants.

## Discussion

Our analysis of copy number variants on the human autosomes has identified an extensive region (∼ 8 MB) on the short arm of chromosome 19, (19p13.3-13.2) that is intolerant of duplication in females, and has shown that there are no other comparable regions in our genome. Most of the genes within this region (approximately 237) show marked skewing from the expected 50: 50 sex ratio of duplications, reflecting paucity of females. The region includes ordinary protein coding genes as well as epigenetic factors – not unexpected because of the large size of some *de novo* amplifications. This also impedes the effort to precisely map the limits of the critical regions.

### Sex differences in variants are attributable to female loss

As sex differences in the incidence of duplications and deletions are not expected, we interpret the skewing to be the result of selective embryonic loss. Female embryos are systematically lost prior to implantation if they duplicate the relevant chromosome 19 genes.

### Sex differences for duplications, but not for deletions support the hypothesis of a single active X

Our observations were exactly as we predicted: We expected that a female with partial trisomy of a gene or genes on either chromosome 1 or 19 might end up with two active X chromosomes, and that this would lead to her demise *in utero*. Conceivably, females with duplications of our candidate region on chromosome 19 are being lost for reasons unrelated to X inactivation; however, in light of our previous studies that implicate chromosome 19, and in absence of similar regions on the other chromosomes, other interpretations of our results seem unlikely.

### Chromosome 19 and candidate genes

Of all human chromosomes, chromosome 19 has the highest density of genes, more than double the genome average [25]. Nearly one-quarter of them belong to tandem-arrayed families. Clearly, most of the genes in our regions of interest are passengers, whereas the drivers have not yet been definitively identified. Table 2 shows some of the more likely driver candidates, as they could repress *XIST*. The DNA methyltransferase *DNMT1* is strategically placed in the middle of the dense19p13.2 zinc finger cluster (Table 3). Although not essential for inactivation of an X, *DNMT1* has been implicated in the process of *Xist* inactivation, previously by Panning and Jaenisch [26]; based on studies of *Dnmt1* deficient mice of both sexes that expressed Xist from their active X’s, these investigators concluded that *Dnmt1* might have a role in inactivating *Xist* on the mouse *active* X. Although primarily functioning as a maintenance DNA methyltransferase, DNMT1 or its splice variant is thought to have a *de novo* methylation function for imprinted genes, including *Xist* [27, 28]. Also in our candidate region, UHRF1, a histone and DNA-binding RING E3 ubiquitin ligase, is an essential co-factor for DNMT1 [21]. And exhibiting some of the greatest skewing are the paired scaffold attachment genes and transcriptional repressors, SAFB and SAFB2, which could assist in mediating *XIST* methylation. All of these factors are functioning in the early mammalian embryo, prior to gastrulation, when random X inactivation occurs [27-30]. Direct testing of candidate genes is impeded by bans on human embryo studies, and the fact that the human embryonic stem cells currently available have already undergone X inactivation. Nonetheless, our observations should stimulate the development of a suitable human assay system.

Our model of an autosomal *XIST* repressor calls for only one dosage sensitive gene. Other drivers may well be dosage insensitive. It is difficult to know how such a dosage sensitive repressor functions, because in normal diploid cells there are two copies of the relevant autosomal repressor and only a single *XIST* locus to silence. It is unlikely that the product of both chromosome 19 genes is needed as deletion of one locus seems to be tolerated, so that the right dosage may require some form of competitive inhibition. Alternatively, physical contact with a single chromosome 19 may be needed to assure that only one *XIST* gene is repressed.

### Potential species differences

There are significant differences in the process of X inactivation among mammals including its timing, presence of parental imprinting and nature of the long non-coding RNAs; such differences are attributable to evolutionary changes within the X inactivation center and the staging of embryogenesis [31]. The clusters of zinc finger genes arose on chromosome 19 after the split of humans from rodents, and reside on chromosome 19 in other primates [22]. In addition, the major region of skewing – on the human chromosome 19 short arm – is found on two different chromosomes in rodents (mouse chromosomes 17 and 9, and rat chromosomes 9 and 8), and the long arm region is found on yet another chromosome. Percec et al., 2003 [32] used ENU chemical mutagenesis to screen for autosomal mutations involved in the initiation of X inactivation in mice. They identified regions of mouse chromosomes 5, 10 and 15 that seemed to affect choice of *inactive* X; none of these chromosomes is orthologous to human chromosome 19. Further identification of the relevant genes will tell us if we share mechanisms with other mammals, or if such species differences reflect the lack of shared strategies to create a single active X.

### Further implications of the loss of human females with partial trisomies

Selective loss of females at the time of X inactivation could help explain why the human male: female ratio at birth is 1.05-1.06 to 1.0 [33]. The exact number differs with country and is influenced by the recent advent of sex selection by prenatal diagnosis; Yet, without doubt more males are born than females throughout the world. The reason for the skewed sex ratio has been enigmatic in absence of bias in gametogenesis or fertilization in favor of Y-bearing sperm, and in light of evidence that more males are lost than females at every stage post-implantation [34, 35]. As X activation is a dosage sensitive process, it is more hazardous for females than for males; males have only a single X to maintain, but females face the danger of activating more than one X chromosome (inactivating more than one *XIST* locus), which would be lethal. It seems that females, who by chance, inherit a duplication of their dosage sensitive *XIST* repressor(s) cannot survive.

The lack of females with partial trisomies of our candidate region on chromosome 19 affirms their selective preimplantation loss. Such a loss of females must contribute to the distorted sex ratio at birth. Just as the absence of autosomal monosomies from studies of fetal wastage tell us which zygotes have been lost, the absence of females with partial trisomies of the region 19: 4.5-12.5 MB and 41-41.5 MB in recognizable pregnancies, documents their preimplantation loss.

### Sex ratios have value in the study of disease

Not only does the sex ratio of copy number variants tell us about fetal loss, but it can also provide insights into disease processes. For many genetic diseases of autosomal origin, including autism and Hirschsprung disease, there is a marked sex difference in the expression of the disease, not apparently attributable to hormones or X-linked genes. We suggest that sex differences in some copy number variants, and not others, may provide useful clues to mechanisms underlying the deviations from expected sex ratios for the risk of disease. Our data draw attention to the potentially significant role played by autosomal copy number variation in establishing the active X, and in mediating gender bias in disease manifestations; therefore they warrant further investigation.

## Materials and Methods

### Search for candidate genes on chromosomes 1 and 19

We searched OMIM and the UCSC Genome Browser for each band along the chromosome to ascertain candidate genes within the region. Such potential candidates for *XIST* repressors included epigenetic factors, histone structural components, histone methyltransferases, DNA methyltransferases, zinc finger proteins, heteronuclear proteins, long non-coding RNAs, other genes implicated in X inactivation and genomic imprinting and undefined open reading frames. They were then subjected to a search in the DECIPHER *database* of all reported variants (duplications, deletions, gains and losses). When a chromosome band was implicated from the DECIPHER search, then all genes within the band, present in that database were assayed. These data are reported in Tables 1, 2 and 3, and S1 Table. Detailed analysis of chromosome 19 was carried out by 500 KB bins from pter to qter, using M: F sex ratios for total copy number variants, duplications and deletions, which were plotted for Fig 3, as described below.

### Ascertainment of sex ratio on all the human autosomes using DECIPHER gains

To systematically sample a sizable fraction of human genes on all the autosomes, we used the available DECIPHER gene list, a compilation of genes with open-access patient sequence variants, or DDD research sequence variants. On September 6^th^ 2016, this list contained 2737 genes, of which 2697 were protein coding. We ordered these genes serially on the chromosomes to identify sex-ratio skewing in regions of interest from chromosome 1 and 19 and then, to determine if other chromosomes showed skewing of similar magnitude to that observed on chromosome 19. As we were interested in any deviation of the expected equal ratio we collected information about the number of total gains (duplications and partial duplications) in both sexes.

### Calculation of posterior rates

Because the lack of gains in either males or females leads to a zero value in either the nominator or denominator of a male/female gain ratio we calculated a posterior rate, which accurately reflects any observed deviation from the expected equal amount of male/female gains=1. The posterior rate was calculated in the following way: (M+a)/(Nm+A)*(Nf+A)/(F+a)) where M is the number of male duplications observed, F is the number of female duplications observed, and Nm and Nf are the number of males and females in the DECIPHER database, and the Dirichlet prior parameters are a=1 and A=2. [36]. For these studies we assumed that the numbers of females and males are approximately equally represented in the DECIPHER database. The posterior sex rate is represented in Figs 2 and 3, S1 Fig and S1 Table. either at specific genes (Fig 2 and S1 Fig, or for total variants, duplications and deletions within 500 KB bins on chromosome 19 (Fig 3 and S1 Table). The approximate band location for each list gene on chromosomes 1 and 19 is represented below each graph in Fig 2 and S1 Fig using ideograms from Idiogram Album by David Adler. (http://www.pathology.washington.edu/research//cytopages/idiograms/human/) Finally, this study makes use of data generated by the DECIPHER community (see acknowledgement).

## Web Resources

The URLs for data presented herein are as follows:

OMIM, https://www.ncbi.nlm.nih.gov/omim

DECIPHER, https://decipher.sanger.ac.uk/

UCSC genome browser, https://genome.ucsc.edu/cgi-bin/hgGateway

## Acknowledgments

We are grateful to many. We thank Patricia A Jacobs for her contributions to, and support of, the single active X hypothesis. Thanks to David Valle for directing us to the DECIPHER database. We also thank Andrew McCallion, Haig Kazazian, Kirby Smith, Hal Dietz, Dimitrios Avramopoulos and Geraldine Seydoux for their helpful comments about the paper, Cate Kiefe for her rendering of Figure 1 and Giovanni Carosso for help with formatting. This study makes use of data generated by the DECIPHER community. A full list of centres that contributed to the generation of the data is available from http://decipher.sanger.ac.uk and via email from. Funding for the DECIPHER project was provided by the Wellcome Trust.

## Supporting Information

**S1 Fig.** (related to Fig 2). Posterior gain rate (M:F) plotted serially on chromosomes 2-18 and 20-22 (Figs A-T) using available genes from DECIPHER. Dashed line depicts equal gain rate.

**S2 Fig.** (related to Fig 3). Permutation testing reveals that skewing in the 19p region is highly significant (p<10^-11^). Calculations show that even after 10^7^ random simulations, it would require 10^4^-10^5^ more simulations to observe such a skewing at least once.

**S1 Table.** (related to Fig 3). Details of data in Fig 3, including sex ratio of total variants, duplications and deletions and number of genes assayed on Chromosome 19 in 500 KB bins.

